# Comparative photosynthetic capacity, respiration rates, and nutrient content of micropropagated and wild-sourced *Sphagnum*

**DOI:** 10.1101/2024.06.06.597854

**Authors:** Anna T. Keightley, Chris D. Field, James G. Rowson, Simon J.M. Caporn

## Abstract

Rapid, effective restoration of degraded peatlands is urgently needed to reduce their current high levels of carbon loss. Re-introduction of *Sphagnum* moss, along with re-wetting, is key to returning carbon sequestration and retention capabilities to northern degraded bogs. Micropropagated (BeadaMoss®) *Sphagnum* has already been applied in large quantities, and more is planned, for restoration projects in Britain and parts of Europe. Comparison with wild-sourced *Sphagnum* material is therefore pertinent to demonstrate its safety and suitability for wide-scale application. Six *Sphagnum* species of micropropagated and wild-sourced origin were studied: *S. capillifolium*, *S. fallax*, *S. palustre*, *S. papillosum*, *S. medium/divinum* and *S. squarrosum*. Micropropagated *Sphagnum* had significantly higher light-saturated photosynthesis (Pmax) rates, little colour expression, an open habit (shoots separated), higher numbers of chloroplasts, and more numerous, smaller shoot apices (capitula) than wild-sourced *Sphagnum*. Potentially, greater numbers of chloroplasts in micropropagated *Sphagnum* facilitate higher photosynthesis rates, driving rapid growth in early-stage plants, particularly in optimum moisture conditions. Pmax rates were associated with lower bulk density (mass/volume) and higher nutrient concentrations in tissues. Micropropagated *Sphagnum*, grown with additional nutrients, showed no sign of nutrient toxicity or limited P or K, despite a N content approaching 30 mg g^-1^ (well above the highest literature-reported concentration threshold), indicating its nutrient-demanding early growth stage. Ranking species according to their Pmax and respiration rates (expressed on a dry matter basis) found that *S. squarrosum* was the highest and *S. medium/divinum* was the lowest, in both micropropagated and wild-sourced *Sphagnum*. Micropropagated *Sphagnum* is similar in form to wild-sourced, can be propagated in large quantities, and due to high photosynthesis rates and absorption of nutrients, is likely to establish well on application to a site where re-wetting has already occurred, therefore making it highly beneficial for restoration of degraded bogs.

## Introduction

*Sphagnum* is instrumental in its role as an ecosystem engineer in creating the cool, wet, acidic, low-nutrient conditions leading to the formation of peat in northern peatbogs, through chemical processes and recalcitrant plant tissues [1–3]. *Sphagnum* absorbs aerially deposited nutrients rapidly and directly into plant tissue [4,5] and stores them efficiently, reducing their availability to the roots of vascular plants [6,7]. Different species have also developed a range of adaptive or phylogenetic traits to specific ecological conditions of light, shade and moisture [8–11], allowing them to outcompete vascular plants in the resulting hostile bog environment [2,7].

Intact peatlands store more atmospheric carbon per hectare than other terrestrial habitats [12,13], however, many peatbogs are degraded through drainage and subsequent land-use change. Greenhouse gas (GHG) emissions from degraded peatlands are substantial, for example in the UK they were estimated at approximately 23.1 Mt CO_2_e yr^-1^ by Evans *et al*. [14] which was 5% of the total GHG emissions for the UK in 2017 [15]. Restoration is vital to reduce this contribution to anthropogenic climate change [16,17]. *Sphagnum* is a key species for bog restoration [18] but the quantities available for re-introduction are insufficient from wild sources in many regions, including the UK, and sites are often protected [19]. BeadaMoss® *Sphagnum* (described below) has been developed to fill that gap in resources and has already proved successful on wide-scale introduction to blanket bogs in the north of England [20,21]. There is an urgent requirement to examine and understand the performance of micro-propagated *Sphagnum* because of the scale of current and future peatland restoration needs.

The *Sphagnum* genus occupies a wide range of peatland ecological niches, and *Sphagnum* species vary widely in their photosynthetic rates, yet the plant and environmental factors driving production and carbon accumulation remain unclear [22]. Loisel *et al*. [23], in a wide-ranging review of global measurements, found *Sphagnum* stem-length growth and photosynthetically active radiation (PAR) were strongly correlated, with PAR a more important growth indicator than moisture levels, albeit for only two species assessed, *S. magellanicum* (probably *S. medium* or *S. divinum:* [24]) and *S. fuscum*. However, studying the same two species, Bengtsson *et al.* [22] found that PAR had only a low annual effect on length increment, and that a range of other factors were more important, although effects were species-specific. These included moisture levels, temperature, Nitrogen deposition and vascular plant cover, with the latter no doubt influencing PAR levels available to *Sphagnum*. Photosynthesis is constrained by moisture levels in bryophytes, as the plants are poikilohydric: too much moisture content limits CO_2_ diffusion and reduces carboxylation, and too low moisture content damages photosynthetic apparatus [25]. Low levels of nutrients, particularly nitrogen, support photosynthesis in mosses, although higher levels can be toxic and promote shading from vascular plants [26,27]. *Sphagna* utilise the same nutrient elements that all plants use for photosynthesis, respiration and growth, but absorb them directly into cells [4] as they have no vascular transportation system for uptake from the soil, and allocate them differently, as nutrient resources are limited in an ombrotrophic bog system [8,28].

Photosynthesis rates generally decline following the ‘successional gradient’ of species in bog development towards ombrotrophic conditions [11]. Species with metabolic strategies such as high bulk density and carotenoid concentration to tolerate drier, unshaded conditions, tend to have reduced rates of growth and photosynthesis [8], and shade-adapted species tend to have high photosynthesis rates [29]. Moreover, Hájek *et al*. [9], in laboratory conditions and under a range of light intensities, found a clear ranking of CO_2_ uptake between species, with those sourced from shaded habitats tending to rank higher than those sourced from open habitats. The authors surmised that species from open habitats suffered persistent photodamage, which reduced photosynthetic capacity, despite photoprotective pigments such as sphagnorubin.

Micropropagation Services Ltd (trading as BeadaMoss®) produces bulk quantities of *Sphagnum* from a few strands of wild-sourced material sourced from bogs in the north of England, primarily the Peak District and Cumbria. Production involves standard tissue-culture techniques of plant division in a sterile, controlled environment, then growing under greenhouse conditions with minimal, bespoke nutrient application [19]. Studying micropropagated *Sphagnum* is an opportunity for novel comparisons of species that have been cultured and grown under optimum light, moisture and nutrient levels, and are at the same stage of development.

Studies into the photosynthetic capacity of different species of *Sphagnum* moss from contrasting sources are pertinent to understanding production and, therefore, carbon sequestration in bogs [23]. Respiration is part of the net CO_2_ exchange, so is also of study interest. The aims of this study were to make a comparison of photosynthesis and respiration rates, examine any differences in chlorocyst (cell containing chloroplasts) size and number of chloroplasts, and differences in nutrient content of six *Sphagnum* species of both tissue-cultured (micropropagated) and wild-sourced plants. A greater understanding of the potential carbon sequestration of each micropropagated species will help direct both product development and restoration efforts where these products are used.

The objectives were, firstly, to measure the CO_2_ uptake (photosynthesis) and emission (respiration) of samples of *S. capillifolium*, *S. fallax, S. medium/divinum, S. palustre, S. papillosum* and *S. squarrosum* in controlled conditions over a range of light intensities. These represent species from a broad environmental range and were readily available in BeadaMoss® greenhouses. Wild-sourced samples were taken from established, naturally occurring colonies in a range of peatland environments. Secondly, micropropagated and wild-sourced samples of the same six species used for photosynthesis rate studies were examined under a microscope and measurements made of chlorocyst size and number of chloroplasts to test for any differences which may influence capacity for photosynthesis. Thirdly, the nutritional content of samples used for photosynthesis measurements were analysed to examine whether levels of nutrients within tissues of micropropagated *Sphagnum*, which is a horticultural product, and wild-sourced *Sphagnum*, had a bearing on their photosynthetic capacity.

Our research questions, which we succeeded in addressing, were:

1) Which *Sphagnum* species, either grown under the same conditions (micropropagated) or grown in the wild, show the greatest photosynthesis and respiration?
2) Is there a difference in photosynthetic capacity and respiration between micropropagated and wild-sourced *Sphagnum* species;
3) Are there differences in chlorocyst size, chloroplast number and nutrient content between micropropagated and wild-sourced *Sphagnum* species which may explain differences in their photosynthetic capacity and respiration;

## Materials and methods

### *Sphagnum* photosynthesis and respiration

Approximately 2 litres of each wild-sourced *Sphagnum* species were collected in August 2017 from ombrotrophic mires and heaths in the north of England and Wales: *S. capillifolium, S. fallax,* and *S. palustre* from Chat Moss Remnants Site of Biological Importance (SBI) (now owned by Lancashire Wildlife Trust, and called ‘Rindle Moss’) (53°27’53.0"N, 2°26’56.4"W), *S. medium/divinum* from Borth Bog (Cors Fochno) (52°30’18.7"N, 4°00’43.5"W) and *S. papillosum* from Ruabon Moor (53°00’11.5"N, 3°08’21.6"W). *S. squarrosum* was collected from Alderley Edge (53°17’52.1"N, 2°12’18.0"W), a wooded basin mire.

Samples of the same species were also collected from BeadaMoss® greenhouses in June-July 2017. Micropropagated *Sphagnum* material is cultivated in glasshouses, where shade is applied in summer and humidity and temperature conditions are controlled, with minimal, bespoke (commercially sensitive information) nutrient application. The samples used were grown from BeadaGel™ (micropropagated *Sphagnum* suspended in a hydrocolloidal gel and applied directly onto the growing-media surface). The species selected were typical of the BeadaMoss® *Sphagnum* grown for restoration work.

All *Sphagnum* samples were acclimatised in a Fitotron growth chamber (Weiss Technik, Lindenstruth-Reiskirchen) for a minimum of 5 days, set to typical summer-time environmental conditions in the local area: 20°C during the day (0600 - 2200 hrs), 12°C during the night, day-time light intensity of 750-800 µmol m^-2^ s^-1^ (values calculated from the nearby Astley Moss Weather Station mean data, 2012 – 2015), 85% humidity. The *Sphagnum* was misted with rainwater as necessary to keep it hydrated.

A literature review was used to determine the range of light levels needed to capture the photosynthetic response of diverse *Sphagnum* species. Haraguchi and Yamada [30] found that optimum light levels for photosynthesis for a range of *Sphagnum* species were between 300 and 500 μmol photons m^-2^ s^-1^ of PPFD (Photosynthetic Photon Flux Density). However, Rice *et al*. [8] found that photoinhibition (to prevent high-light damage) in *Sphagnum* occurs at 800 μmol photons m^-2^ s^-1^ and Loisel *et al*. [23] reported an optimum of 500 to 900 μmol photons m^-2^ s^-1^. Hájek *et al*. [9] found that the light saturation point for all *Sphagna* they studied was similar at an average of 2124 ± 86 (SE) µmol photons m^-2^ s^-1^, much higher than for other studies. In this study, 800 μmol photons m^-2^ s^-1^ was the highest light level.

Five replicate samples of each species from each type (micropropagated and wild-sourced) were cut to 3 cm lengths from the 2-litre bulk amounts of *Sphagnum* at growing shoot density and placed in a 5 cm diameter x 3 cm high clear acrylic cylinder (i.e., not a standard number of shoots, or compressed to fit) fitted with a mesh base to allow air circulation through the samples (Fig 1), with the top surface of the sample level with the top of the cylinder. An LGR™, Ultraportable Greenhouse Gas Analyser (UGGA), Model 915-0011 (Los Gatos, Research, Palo Alto, CA, USA) (LGR) was fitted to a 500 ml sealable clear-glass chamber via tubing through air-tight ports in the lid, and *Sphagnum* samples placed in the chamber for analysis. Change in CO_2_ concentration within the chamber was measured over 2 minutes and a light response curve determined for each species starting from light intensities of 800 µmol photons m^-2^ s^-1^ to zero in increments of 50 µmol photons m^-2^ s^-1^ for each sample. A Skye PAR quantum sensor, integral to the growth chamber, was placed on a level with the sample being assessed to set an accurate light intensity. The reduction in light transmission through the chamber (8%) was accounted for by increasing the light intensity in the cabinet accordingly for each light intensity measurement. Samples were lightly misted with rainwater after each measurement and adapted to each change of light level between measurements for approximately 5 minutes.

**Fig 1.**
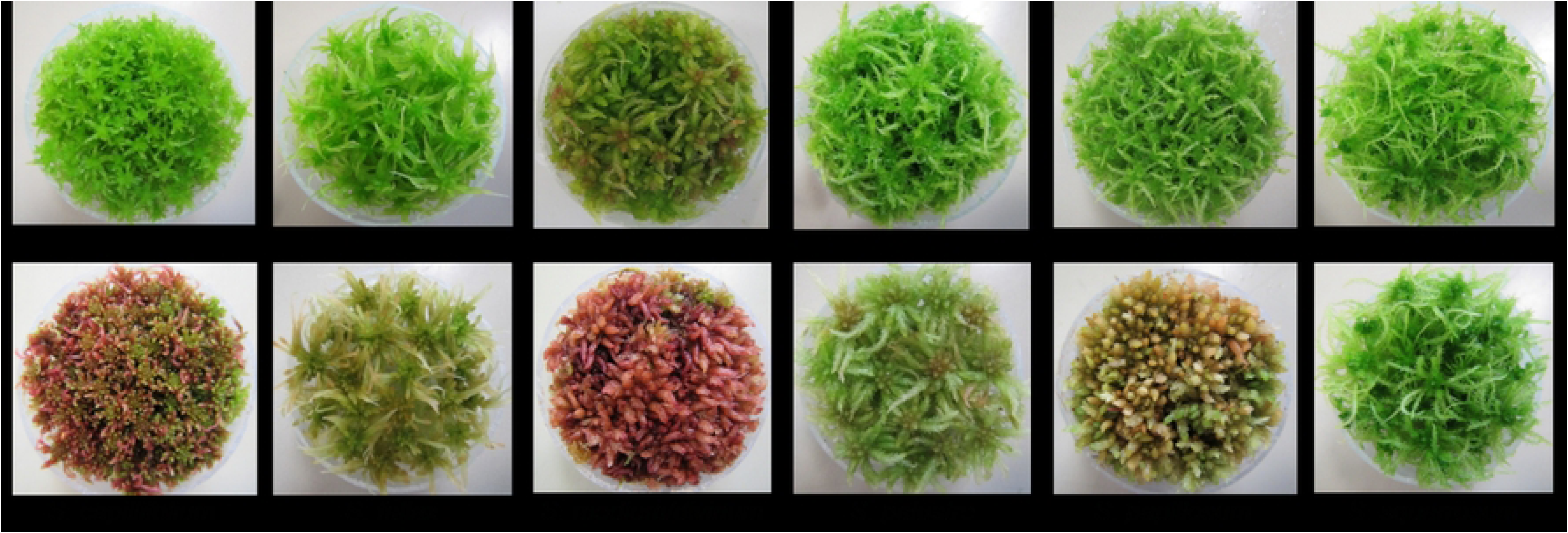
Examples of *Sphagnum* species samples for analysis of photosynthesis rate. Micropropagated (top row of pair) and wild-sourced *Sphagnum* species samples show typical visual differences in pigmentation and form.

The samples were photographed prior to measurements (Fig 1) and the number of capitula per sample counted. Samples were weighed to check for water loss between measurements, and the chamber was closed between each light level measurement to reduce drying. The LGR flow rate was 0.8 litres min^-1^ with space between gas inlet and outlet points; air was released into the chamber along a small pipe with holes along the length to encourage mixing (LGR low flow rate threshold for analysis is ∼0.35 litres min^-1^ [personal communication, Lewis John, LGR™ Sales Representative]).

Net photosynthesis or respiration rate was calculated through Microsoft Excel from the rate of CO_2_ depletion or increase using linear regression over a two-minute period, and further expressed by surface area of the plant chamber (As) and subsequently by total plant dry weight (DW). Values are expressed using the leaf gas exchange sign convention whereby plant uptake of CO_2_ from the atmosphere is expressed as positive and loss to the atmosphere is expressed as negative. The calculation to determine the photosynthesis (CO_2_ uptake) or respiration (CO_2_ emission) rate (adapted from Dossa *et al.* [31]) is:

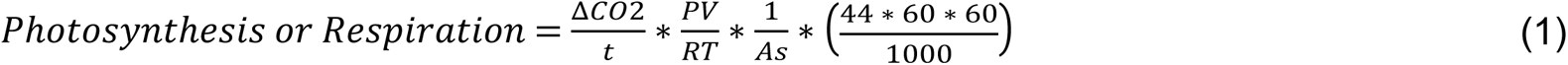

(P (atm) = atmospheric pressure; V (m^3^) = chamber volume; R (L atm mol^-1^ K) = universal gas constant; T (K) = gas temperature in Kelvin; As (m^2^) = sample surface area; 44 g mol^-1^ = molecular weight of CO_2_); Photosynthesis or Respiration = g CO_2_ m^-2^ h^-1^.

### *Sphagnum* samples bulk density

High shoot density and bulk density are physiological aspects of *Sphagnum’*s protective adaptation to open conditions and may lead to reduced photosynthesis rates, so bulk density measurements were made to quantify this. After photosynthesis measurements, samples were dried overnight at 105°C (temperature used by other researchers, e.g., Limpens and Berendse [32], McNeil and Waddington [33]) to obtain dry weight for calculations and for nutrient analysis. Dry weight bulk density was calculated from the dry weight divided by the known sample volume.

### *Sphagnum* samples nutrient analysis

Nitrogen content of the samples was obtained using a LECO FP628 elemental analyser. A minimum of 0.05 g of dry *Sphagnum* was used per sample. Each sample measured for photosynthesis (*n* = 60) was analysed, plus 4 replicates of one species from each type (micropropagated and wild-sourced) to determine experimental error (total *n* = 68).

For other elements, samples (as above) were prepared for Inductively Coupled Plasma Optical Emission spectroscopy (ICP-OES) through acid and microwave digest, using HNO3 S.G. 1.42 (> 68%) PrimerPlus-Trace analysis grade in a CEM Mars Xpress 5 Microwave, and the diluted solution was analysed through ICP-OES (Thermo Fisher Scientific, Waltham, US).

#### 2.4 *Sphagnum* cell measurements and analysis

In addition to *Sphagnum* measured for photosynthesis and nutrient analysis, small samples (several strands) of *Sphagnum* for microscopic analysis were collected in September 2019 from BeadaMoss® unheated greenhouses (daytime temperature ∼20°C) and from natural sources and assessed over the following week. Wild-sourced *Sphagnum* was from sites in the north of England: *S. fallax*, *S. medium/divinum* and *S. palustre* from Cadishead Moss (53°27’07.9"N, 2°27’18.9"W); *S. capillifolium* and *S. papillosum* from Astley Moss (53°28’32.2"N, 2°27’15.5"W), and *S. squarrosum* from Windy Bank Wood (53°28’15.3"N, 2°28’59.6"W). Leaves on divergent branches just below the capitula are generally used for microscopic observation of *Sphagnum* [34,35]. The cell structure changes across the leaf from proximal to distal ends and from edge to centre, and between convex and concave aspects [35,36]. For standardisation of results in this study, leaves from branches just below and immediately surrounding the capitulum (as indicated in Fig 2) were observed and measurements made centrally, on the concave aspect.

**Fig 2.**
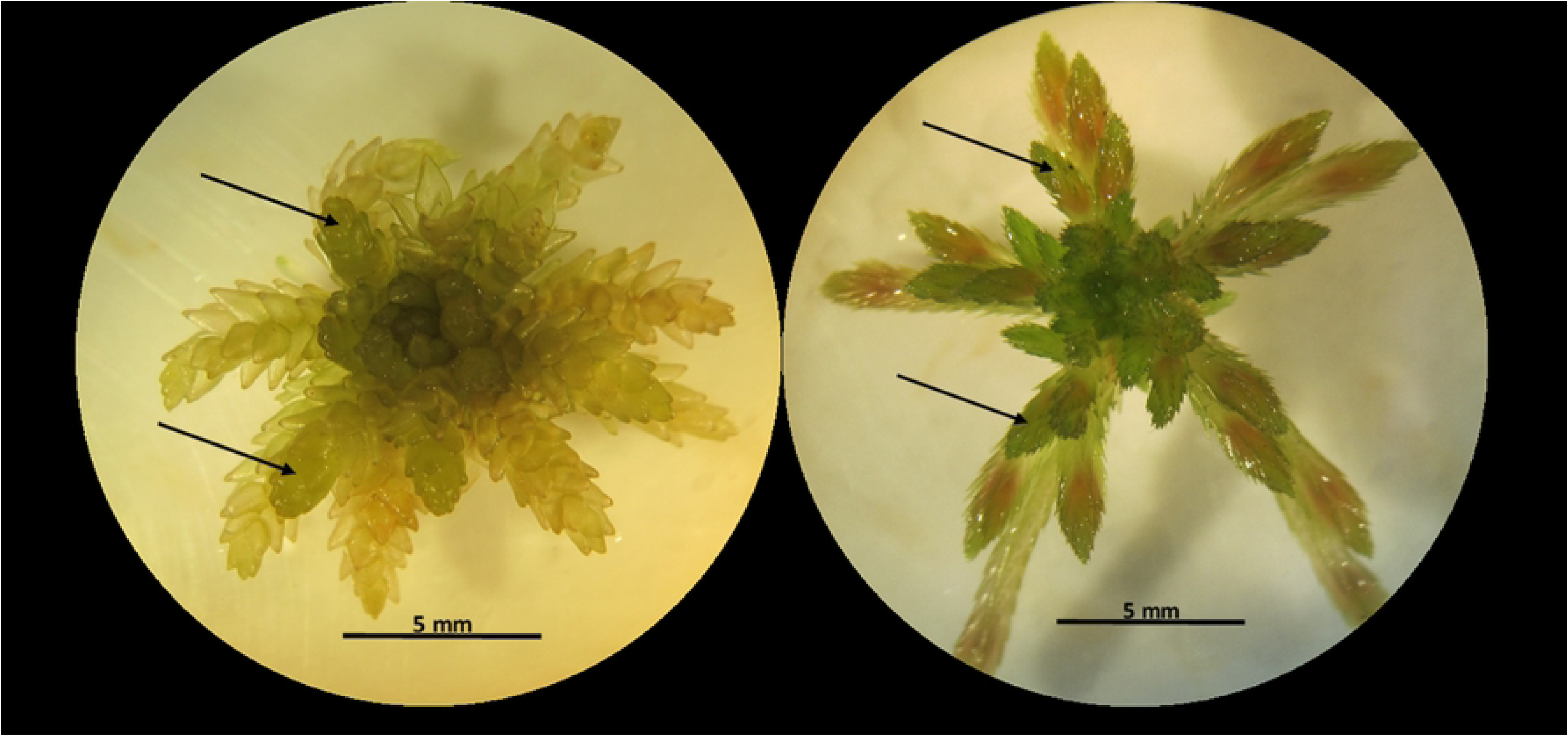
Examples of capitula used for microscopic study; arrows indicate typical branch selection.

Leaves from three branches from below three capitula of each sample (i.e., 9 leaves) were removed onto a slide and photographed using a Brunel Eyecam Plus attached to a compound microscope at 1000X magnification; cell dimensions were measured after calibration at the same magnification. The collection of chlorocysts (cells containing chloroplasts) surrounding a hyalocyst (a dead, thin-walled and hollow cell with a water storage function) are generally made up of six segments of varying size, which are sometimes subdivided. The width (rather than length, because of sub-division of some cells) of six appropriate segments was measured centrally (Fig 3), and the number of chloroplasts counted in each. A mean value was calculated from each set of measurements for each leaf, and thus there were 9 values for chlorocyst width and for number of chloroplasts for each species. Where a chlorocyst was subdivided it was treated as a single entity, and only chloroplasts in the chlorocysts immediately surrounding a hyalocyst were included in measurements.

**Fig 3.**
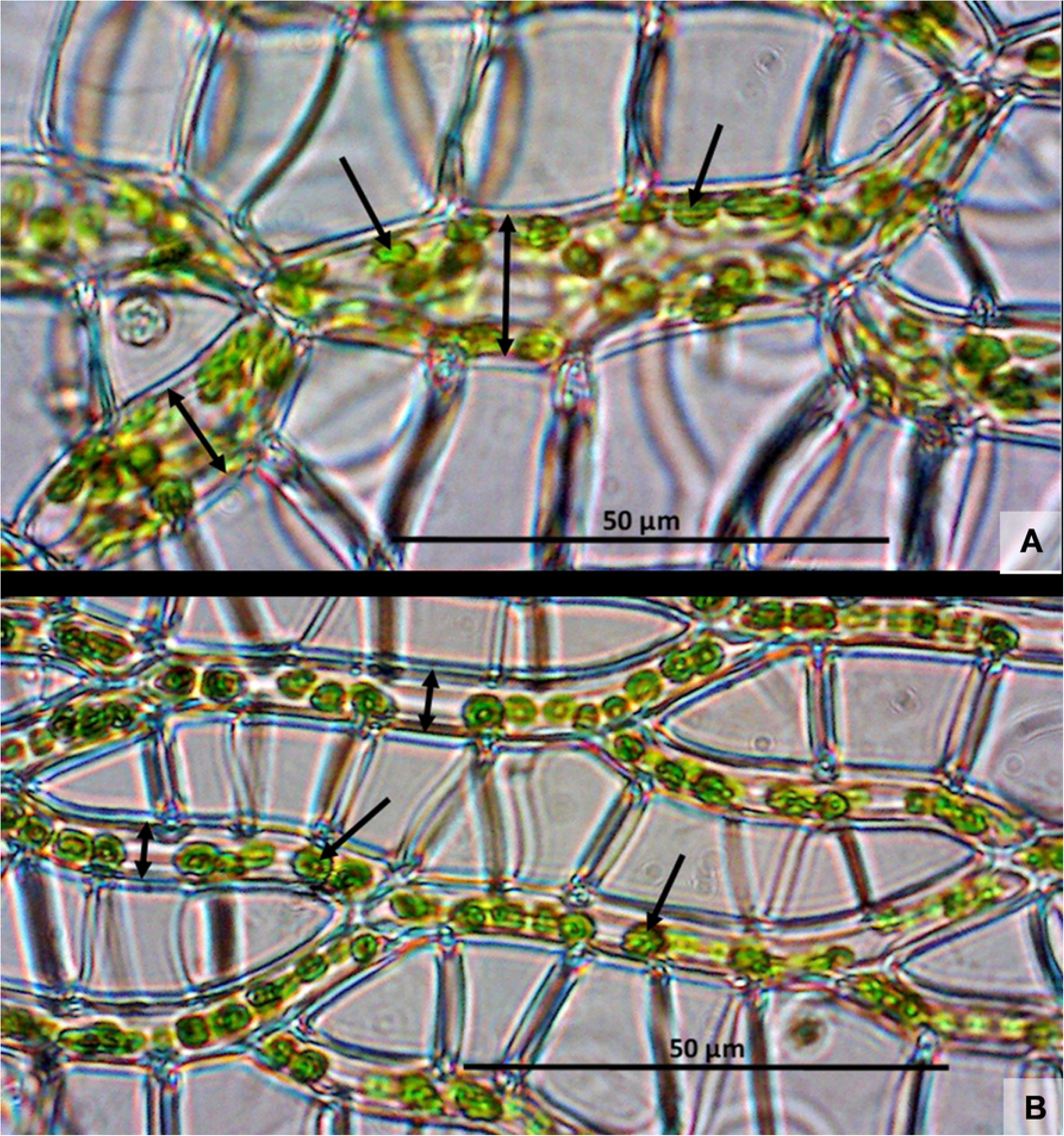
Examples of chlorocyst width measurement locations (double-ended arrows) and chloroplasts (single-arrowed) (A: *S. palustre* and B: *S. squarrosum*).

#### 2.5 Data analysis

Data were analysed using IBM SPSS Statistics for Windows (Version 25.0. Armonk, NY: IBM Corp.) and PAST: Paleontological Statistics Software Package for Education and Data Analysis [37] where indicated. Data were tested for normality using Shapiro Wilk tests. Data for maximum photosynthesis (P_max_), respiration rates, number of capitula per sample, fresh (FW) and dry weight (DW) bulk density and also data for microscopic measurements of chlorocyst width and number of chloroplasts, were found to be normally distributed, and so parametric t-tests were used to test differences between *Sphagnum* types (micropropagated and wild-sourced), and one- and two-way ANOVA with post-hoc Tukey HSD to test differences between species within types, and between types for each species. Dependency of respiration rates on P_max_ rates was tested using linear regression. Data for nutrients were not normally distributed, and so non-parametric independent variable tests (Mann-Whitney U) were used to test differences in distribution between *Sphagnum* types (micropropagated and wild-sourced). Associations between nutrient levels, P_max_ and respiration rates, and *Sphagnum* type and species, were examined with a correlation matrix through Principal Component Analysis using PAST software [37]. Statistical significance was determined with a *p*-value <0.05.

## Results

### *Sphagnum* photosynthesis and respiration

The response of net Photosynthesis rate (P_n_) to changing light levels showed a similar pattern across all species in both micropropagated and wild-sourced samples (Fig 4), reaching a maximum P_n_ (P_max_) between 400 and 650, and 400 and 750 µmol m^-2^ s^-1^ respectively (Table 1) and either levelling or reducing thereafter. Significant inter-species differences were seen within both micropropagated and wild-sourced *Sphagnum* types for P_max_ (*F* = 11.29 and 115.9 respectively; *p* < 0.001, *df* = 5 for both) and Respiration rates (*F* = 12.0 and 39.15 respectively; *p* < 0.001, *df* = 5 for both). Significant differences between species are indicated on Fig 5. P_max_ was significantly higher in micropropagated than in wild-sourced samples overall (*t* = 8.647, *p* < 0.001, *df* = 58). P_max_ of each species was higher in micropropagated than wild-sourced types (significant differences between species from two-way ANOVA Tukey post-hoc tests are indicated in Table 1). The P_max_ rates across micropropagated samples were less variable than wild-sourced samples (coefficient of variation of 32.2% and 74.7% respectively). Respiration rates were significantly greater in micropropagated than in wild-sourced samples overall (*t* = 5.816, *p* < 0.001, *df* = 58), and by species (not significantly for *S. fallax* or *S. palustre*).

**Fig 4.**
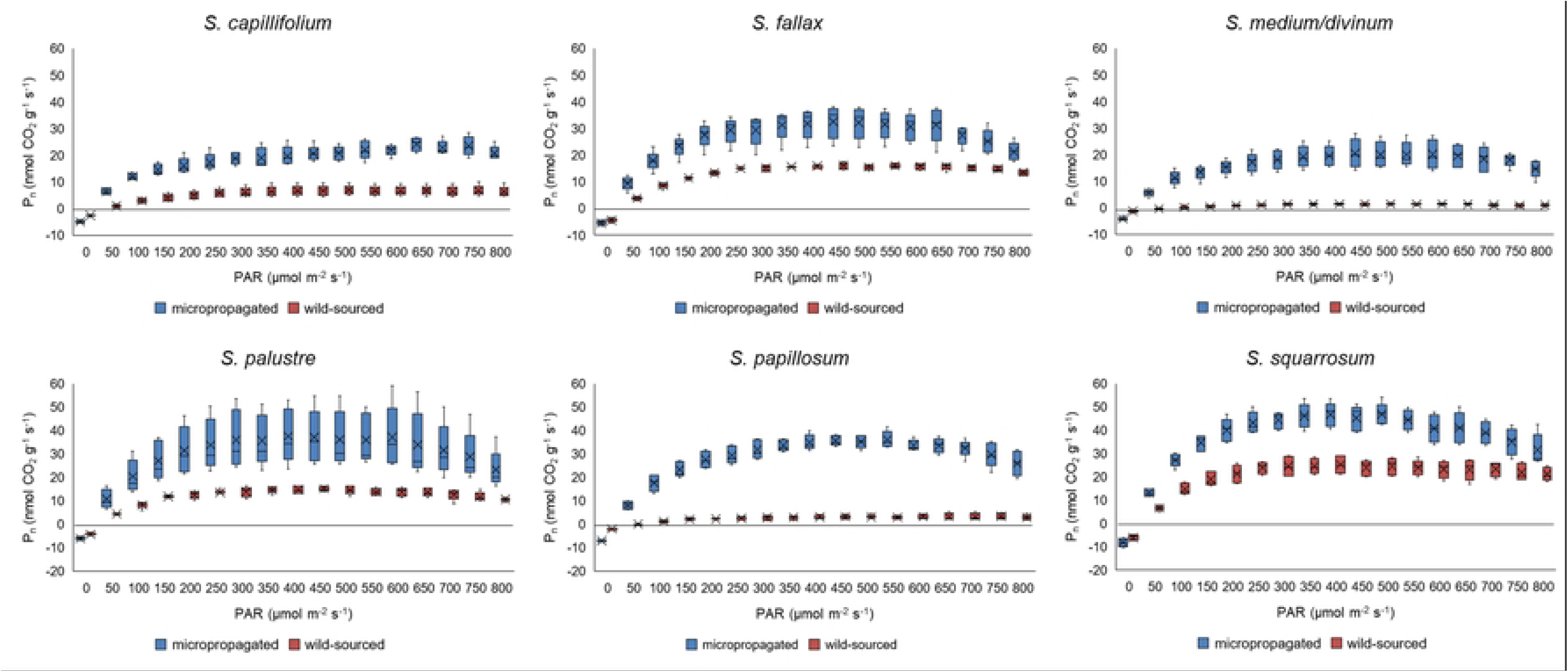
Comparison of micropropagated and wild-sourced *Sphagnum* species net photosynthesis (P_n_) response to light.

**Fig 5.**
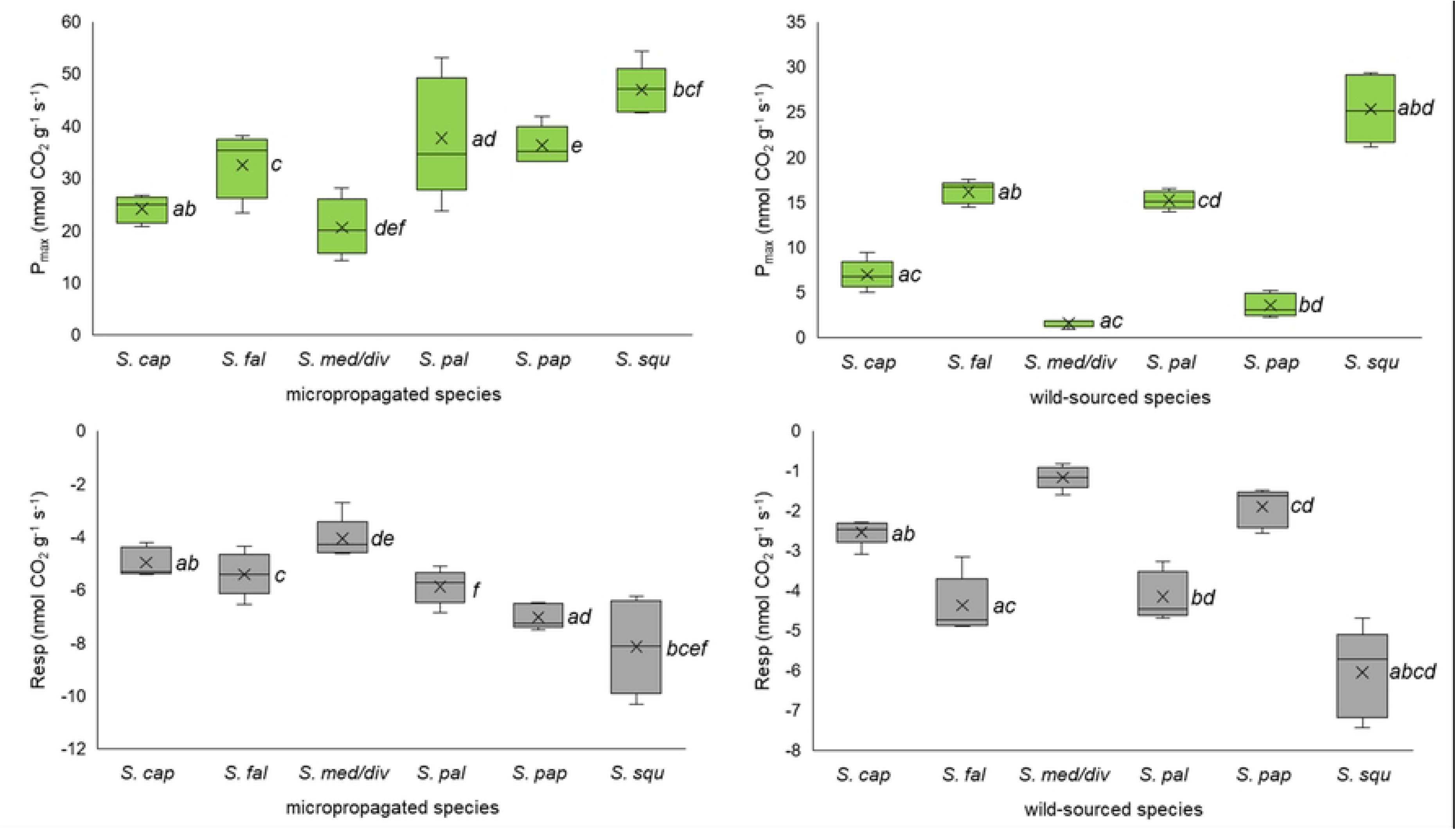
Inter-species differences in light-saturated photosynthesis (P_max_) and respiration (Resp) rates within *Sphagnum* types.

**Table 1.** Comparison of light-saturated photosynthesis (with associated PAR level) and respiration rates of *Sphagnum* species within each type.

| *Sphagnum*<br>species | micropropagated |  |  | wild-sourced |  |  |
| --- | --- | --- | --- | --- | --- | --- |
| | PAR | $P_{max}$ | Respiration | PAR | $P_{max}$ | Respiration |
| *S. squarrosum* | 500 | 46.94 ± 4.74 ** | -8.15 ± 1.77 * | 400 | 25.32 ± 3.75 | -6.05 ± 1.11 |
| *S. palustre* | 400 | 37.69 ± 11.58 ** | -5.88 ± 0.66 | 450 | 15.23 ± 1.00 | -4.15 ± 0.60 |
| *S. papillosum* | 550 | 36.28 ± 3.65 ** | -7.03 ± 0.47 ** | 750 | 3.55 ± 1.27 | -1.91 ± 0.49 |
| *S. fallax* | 450 | 32.54 ± 6.17 ** | -5.42 ± 0.82 | 450 | 16.14 ± 1.25 | -4.37 ± 0.72 |
| *S. capillifolium* | 650 | 24.12 ± 2.53 ** | -4.97 ± 0.55 * | 500 | 6.97 ± 1.64 | -2.54 ± 0.31 |
| *S. medium/divinum* | 450 | 20.57 ± 5.51 ** | -4.07 ± 0.78 ** | 550 | 1.59 ± 0.42 | -1.17 ± 0.29 |

Light-saturated Photosynthesis ( $P_{\max}$ ) with associated Photosynthetically Active Radiation (PAR $\mu\text{mol m}^{-2} \text{s}^{-1}$ ) level and respiration rates of samples, expressed by dry weight ( $\text{nmol CO}_2 \text{g}^{-1} \text{s}^{-1}$ ) ordered greatest to least $P_{\max}$ by micropropagated species, paired with wild-sourced equivalents. Leaf gas exchange sign convention used (i.e., $\text{CO}_2$ uptake positive and $\text{CO}_2$ emission negative). Values are mean ( $n = 5$ ) ± SD. Significant differences (two-way ANOVA Tukey post-hoc tests) between each pair are indicated: \* $p < 0.05$ ; \*\* $p < 0.001$

A light intensity range from 0 to 800 µmol m^-2^ s^-1^ was used to test each sample. Results are shown on a dry weight basis (*n* = 5). Crosses indicate the mean, lines indicate the median, and interquartile range is exclusive with maximum and minimum values that aren’t outliers indicated by the whiskers.

Light-saturated Photosynthesis (P_max_) with associated Photosynthetically Active Radiation (PAR µmol m^-2^ s^-1^) level and respiration rates of samples, expressed by dry weight (nmol CO_2_ g^-1^ s^-1^) ordered greatest to least P_max_ by micropropagated species, paired with wild-sourced equivalents. Leaf gas exchange sign convention used (i.e., CO_2_ uptake positive and CO_2_ emission negative). Values are mean (*n* = 5) ± SD. Significant differences (two-way ANOVA Tukey post-hoc tests) between each pair are indicated: \**p* < 0.05; \*\**p* < 0.001 Statistically significant differences between species on each graph are indicated by shared letters.

Micropropagated species are ranked from highest to lowest P_max_ in Table 1, but the ranking for wild-sourced species was ordered differently for both P_max_ and respiration rates. However, ranking of micropropagated and wild-sourced species with the highest (*S. squarrosum*) and lowest (*S. medium/divinum*) P_max_ and respiration rates were the same.

The ratio of P_max_ to respiration on a DW basis was consistently higher in micropropagated samples (5.58) than wild-sourced samples (3.41). However, there was a strong positive linear regression between P_max_ and respiration rates in wild-sourced samples (R^2^ = 0.90, *p* < 0.001) which was not as evident in micropropagated samples (R^2^ = 0.48, *p* < 0.001).

Weight (moisture) loss from samples during assessment through the range of light intensities was 6.9 ± 2.4% and 7.4 ± 2.9% for micropropagated and wild-sourced respectively; minimum and maximum were 4.4% (*S. medium/divinum*) and 10.5% (*S. squarrosum*) in micropropagated samples and 4.6% (*S. papillosum*) and 9.7% (*S. fallax*) in wild-sourced samples. Moisture content of samples at P_max_ ([Sample P_max_ fresh weight - sample dry weight] / sample dry weight x 100) was 2335 ± 420% (CV = 18%) (micropropagated) and 1551 ± 320% (CV = 21%) (wild-sourced).

There were significantly more capitula per sample in micropropagated than in wild-sourced *Sphagnum* overall (mean ± SD = 41.8 ± 24.2 and 27.0 ± 18.7 respectively; *t* = 2.635, *p* = 0.011, *df* = 58) (Examples in Fig 1). As noted in the Methods, samples were placed carefully in a cylinder for analysis at growing shoot density. Moreover, there were more capitula in micropropagated than in wild-sourced samples of each species, although (on two-way ANOVA post-hoc Tukey HSD) this was not significant for *S. fallax*, *S. medium/divinum* and *S. squarrosum* (*S. capillifolium* and *S. papillosum p* < 0.001; *S. palustre*: *p* < 0.01, *n* = 10).

## Bulk Density

The FW and DW bulk density were greater in wild-sourced than in micropropagated samples (not *S. fallax* or *S. palustre* by FW) (*F* = 22.3 [FW], *F* = 73.6 [DW]; *p* < 0.001, *df* = 11 for both) (Fig 6). Differences between types (micropropagated and wild-sourced) were significant (ANOVA post-hoc Tukey HSD) for *S. capillifolium* (FW *p* = 0.019, DW *p* < 0.001) *S. medium/divinum* (FW *p* = 0.003, DW *p* < 0.001) and *S. papillosum* (FW and DW *p* < 0.001) [*n* = 10 throughout]. Within types, the bulk density across micropropagated samples was less variable than wild-sourced samples (coefficient of variation of 34.7% and 45.3% by FW and 25.0% and 44.5% by DW respectively) with greater bulk density in *S. capillifolium, S. medium/divinum* and *S. papillosum* than other species in wild-sourced samples. There was a statistically significant difference between species within both micropropagated and wild-sourced samples as determined by one-way ANOVA (FW: *F* = 4.71, *p* = 0.004, *F* = 44.65, *p* < 0.001 and DW: *F* = 3.99, *p* = 0.009, *F* = 70.37, *p* < 0.001 respectively). One-way ANOVA post-hoc Tukey HSD statistically significant differences are shown on Fig 6.

**Fig 6.**
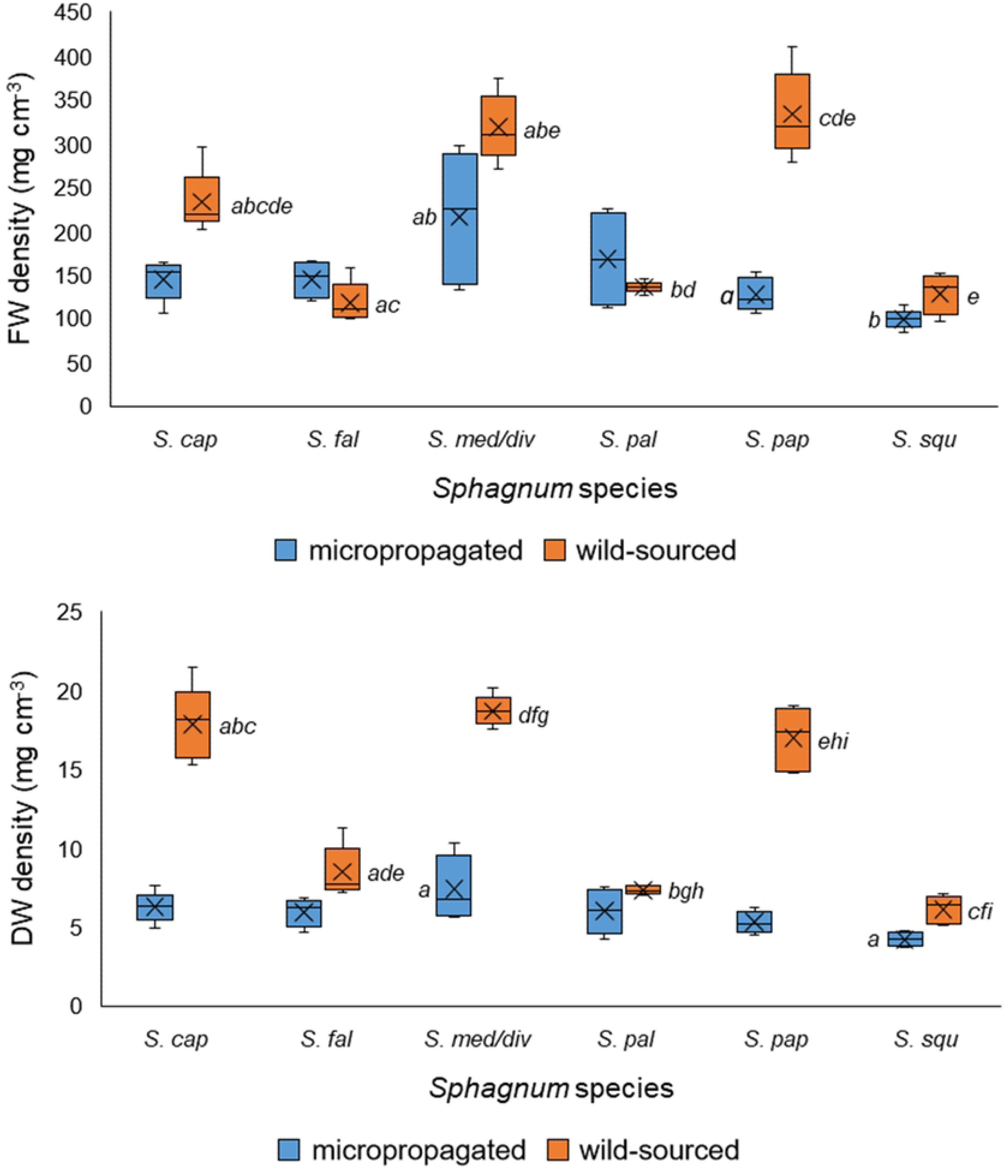
Comparison of micropropagated and wild-sourced *Sphagnum* density by fresh weight (FW) and dry weight (DW). Crosses indicate the mean, lines indicate the median, and interquartile range is exclusive. Shared letters within types (micropropagated and wild-sourced) indicate significant differences in density between species (one-way ANOVA post-hoc Tukey HSD).

Linear regression revealed significant negative relationships between DW bulk density (mass/volume) and both P_max_ (DW) and respiration (DW) (P_max_ and respiration rates decreased as bulk density increased) of micropropagated and particularly wild-sourced samples (P_max_: R^2^ = 0.573 and 0.827 respectively; respiration: R^2^ = 0.503 and 0.789 respectively; *p* < 0.001, *df* = 29 throughout).

## Nutrient content

Macronutrient (Ca, K, Mg, N, P, S) concentrations were significantly higher in micropropagated than in wild-sourced samples overall (Mann-Whitney U test: S: *p* = 0.001, all other elements: *p* < 0.001; *n* = 60 throughout) (Table 2). N made up the largest proportion of the macronutrient content in all species throughout, and there was a high proportion of K in all micropropagated samples and in most of the wild-sourced samples. In the Principal Components Analysis (Fig 7), there was a noticeable clustering of wild-sourced samples away from micropropagated *Sphagnum*, and a disassociation from increasing macronutrients (Ca, Mg, P, N, K, S) in wild-sourced samples (Fig 7A) apart from *S. squarrosum* (a minerotrophic species), which was more closely associated with micropropagated *Sphagnum*. In contrast, micropropagated *Sphagnum* was associated with macronutrients, although there were differences between macronutrients and species. N, P and K were closely associated, and there was an association between these macronutrients and micropropagated *S. palustre, S. papillosum and S. squarrosum*.

**Fig 7.**
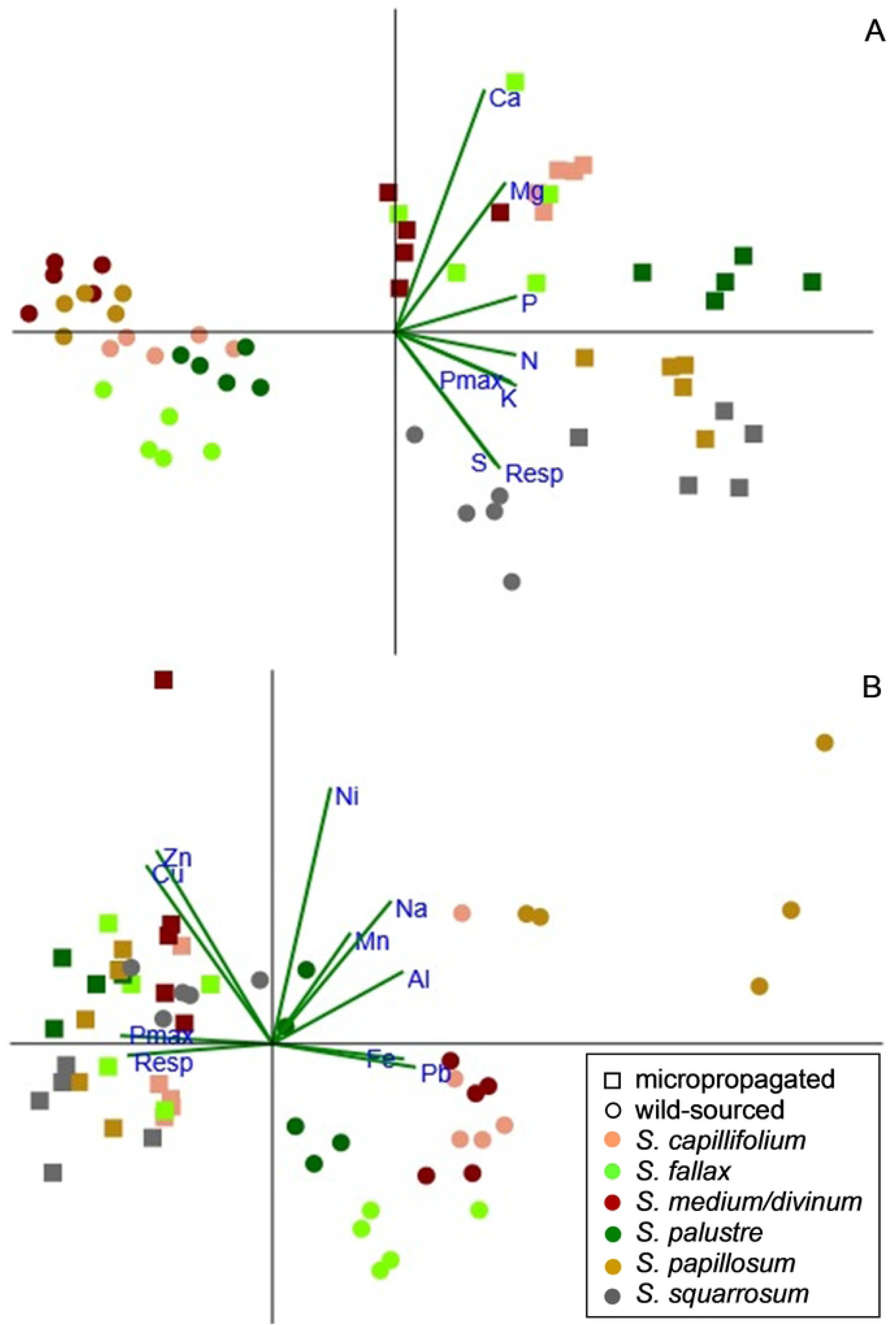
Principal component analysis (PCA: correlation matrix) of Sphagnum element content. PCA graphs show A: macronutrient and B: micronutrient and trace element content of *Sphagnum* samples with P_max_ and respiration by dry weight.

**Table 2.**
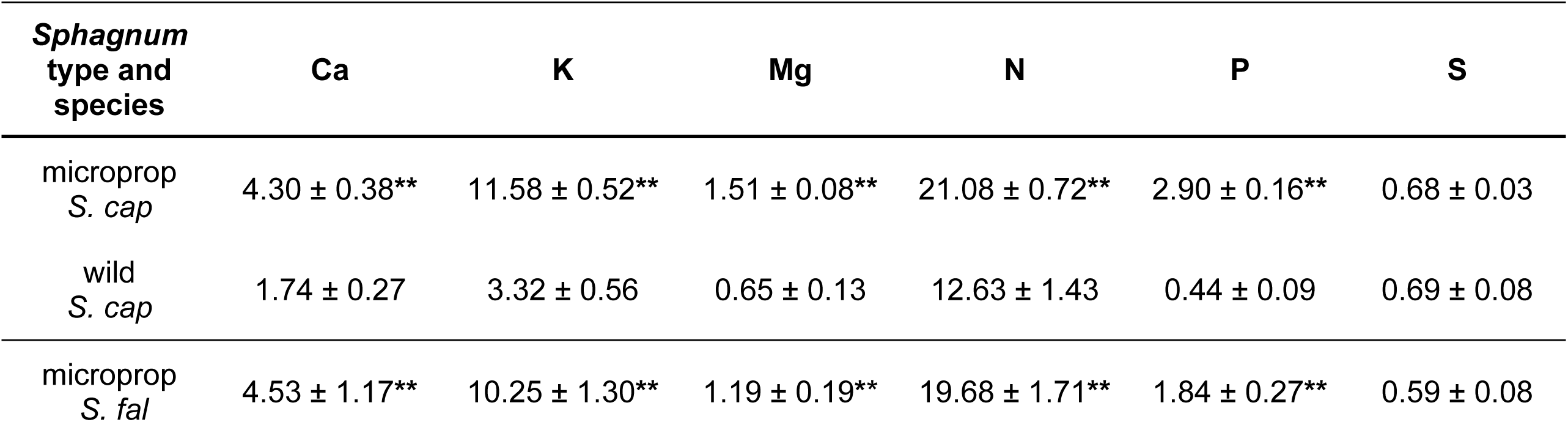
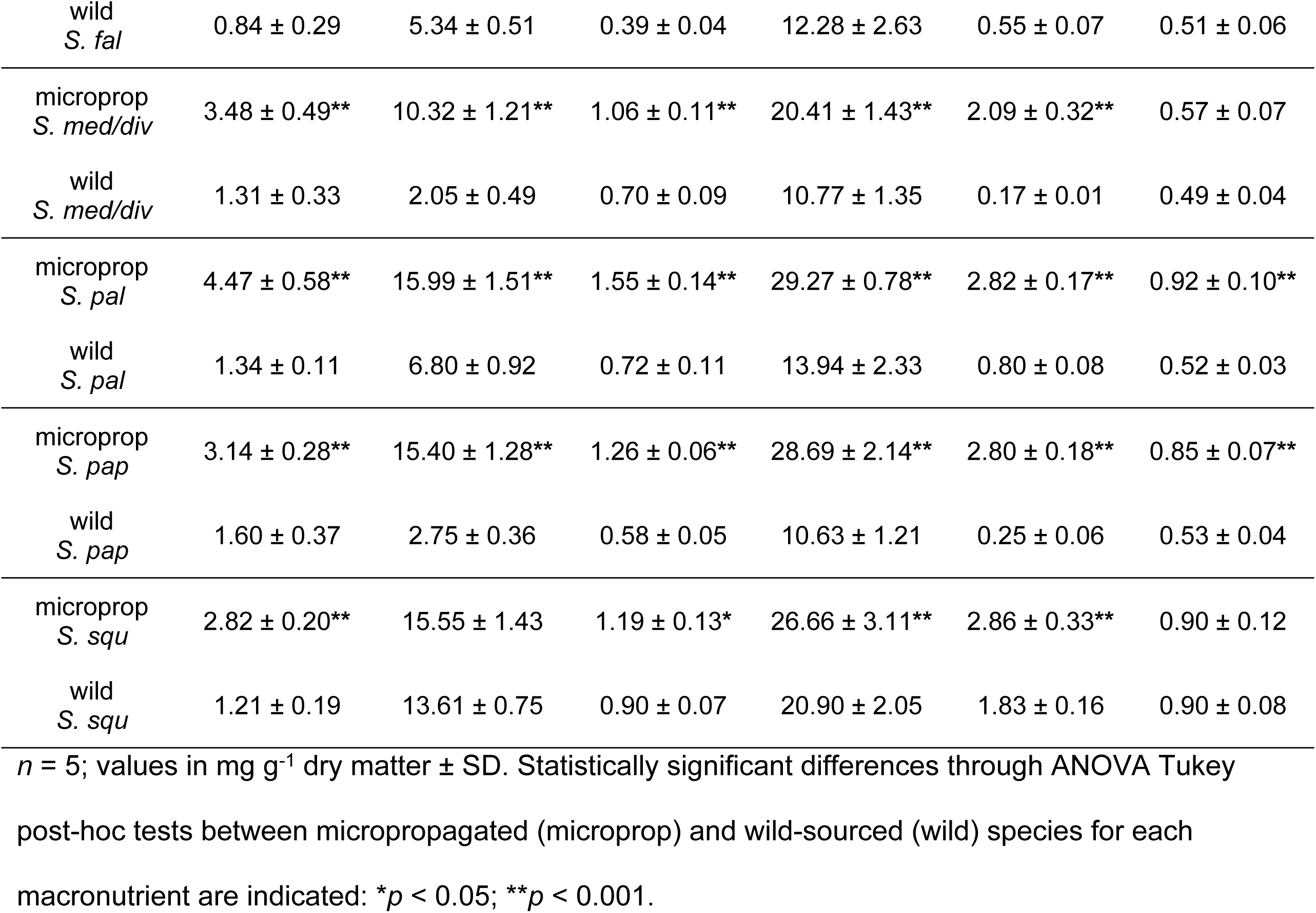
Macronutrient content of *Sphagnum* samples.

Levels of trace elements Al, Fe, Mn, Na, Ni and Pb were higher in wild-sourced than micropropagated samples overall (Al, Fe, Pb: *p* < 0.001; Mn: *p* = 0.01; Na and Ni: *p* NS; *n* = 60 throughout) (Table 3) with the highest levels of Al, Fe, Mn and Na in *S. papillosum*. Levels of micronutrients Cu and Zn were higher in micropropagated than wild-sourced samples overall (*p* < 0.001; *n* = 60) (although not in *S. squarrosum*). Na made up the largest proportion of the micronutrient and trace element content in all samples. In Principal Components Analysis (Fig 7B) there was a clustering and association with Cu and Zn in micropropagated samples together with wild-sourced *S. squarrosum*. Other samples of wild-sourced *Sphagnum* were only loosely grouped by species and more closely associated with micronutrients other than Cu and Zn. Cu and Zn were closely associated, as were Fe and Pb, and Mn and Na. *S. papillosum* appeared to have a strong association with Na and Al, and *S. capillifolium, S. medium/divinum* and *S. fallax* with Fe and Pb.

**Table 3.**
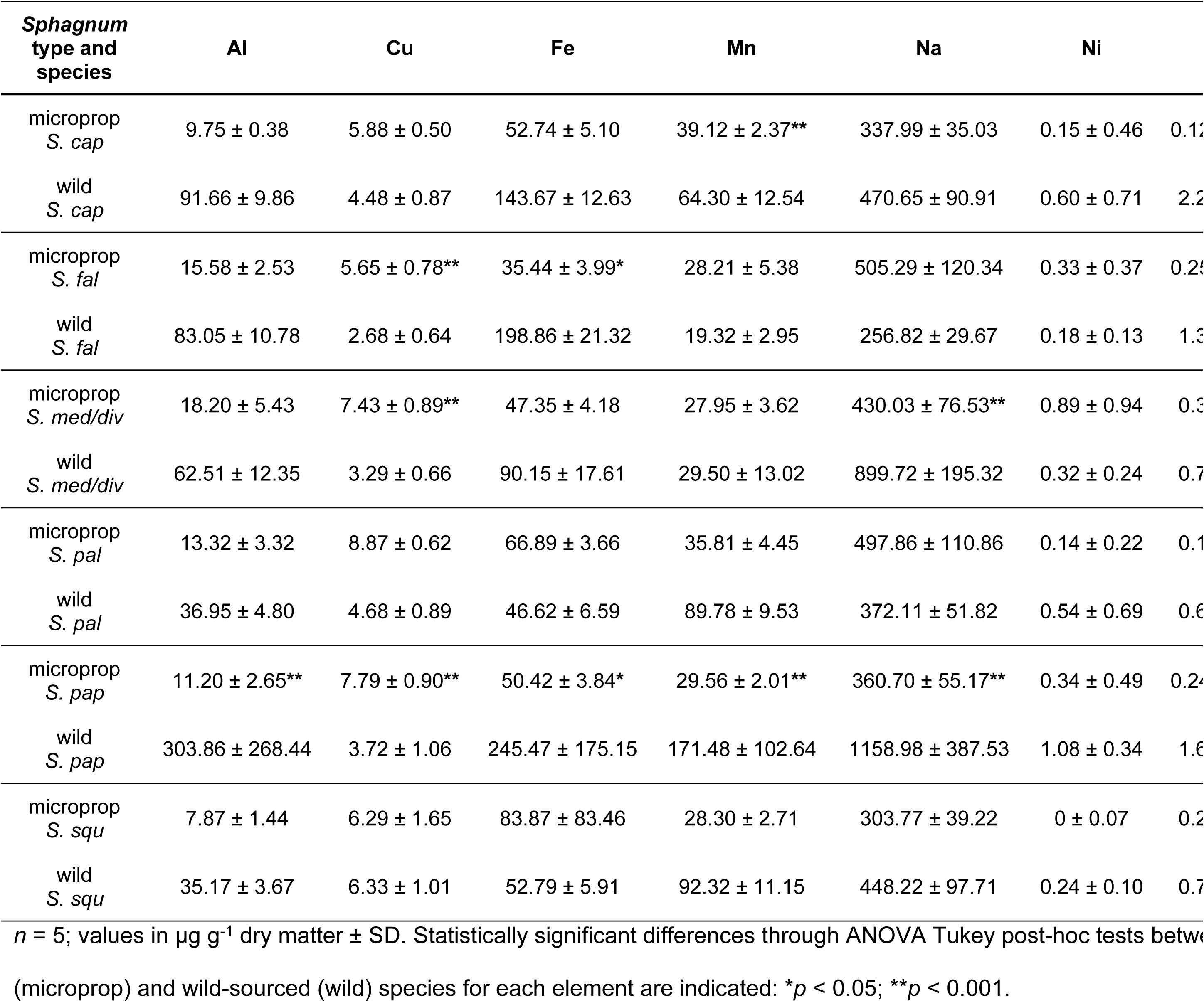
Micronutrient and trace element content of *Sphagnum* samples.

There was a lower N:P and N:K ratio, and a lower variation between species in micropropagated than in wild-sourced samples: N:P = 9.61 ± 1.24 (CV = 12.9%) and 31.18 ± 19.54 (CV = 62.7%) respectively, and N:K = 1.85 ± 0.09 (CV = 4.9%) and 3.13 ± 1.40 (CV = 44.8%) respectively. The N:P and N:K ratios in wild-sourced *S. squarrosum* (11.41 and 1.54), and to a lesser extent, *S. palustre* (17.51 and 2.05) were the most similar to their micropropagated equivalents, and those of wild-sourced *S. medium/divinum* (64.59 and 5.24) and *S. papillosum* (42.56 and 3.86) the most dissimilar.

## Microscopic analysis

There were significant differences between the groups of species by type (micropropagated or wild-sourced) in both chlorocyst width and number of chloroplasts (*F* = 49.9, *F* = 33.7 respectively; *p* < 0.001, *df* = 11 for both) (Table 4). There were no significant differences (ANOVA post-hoc Tukey HSD) in chlorocyst width between micropropagated and wild-sourced species individually (Table 4) except for *S. squarrosum* (*p* < 0.05 wild-sourced > micropropagated). The widest chlorocysts were recorded in *S. palustre* (micropropagated and wild-sourced samples). There were more chloroplasts in micropropagated compared to wild-sourced species in all but *S. squarrosum* (Table 4) and differences were significant (ANOVA post-hoc Tukey HSD) for *S. capillifolium and S. palustre (p* < 0.05) and *S. papillosum (p* < 0.001). The greatest chloroplast numbers in each type were found in *S. palustre* and *S. papillosum* (micropropagated) and *S. palustre* (wild-sourced). Physical differences between micropropagated and wild-sourced samples are limited to reduced colour expression (*S. capillifolium* and *S. medium/divinum*) (Fig 1) and maturity (*S. papillosum* cell papillae) in micropropagated *Sphagnum* (Fig 8).

**Fig 8.**
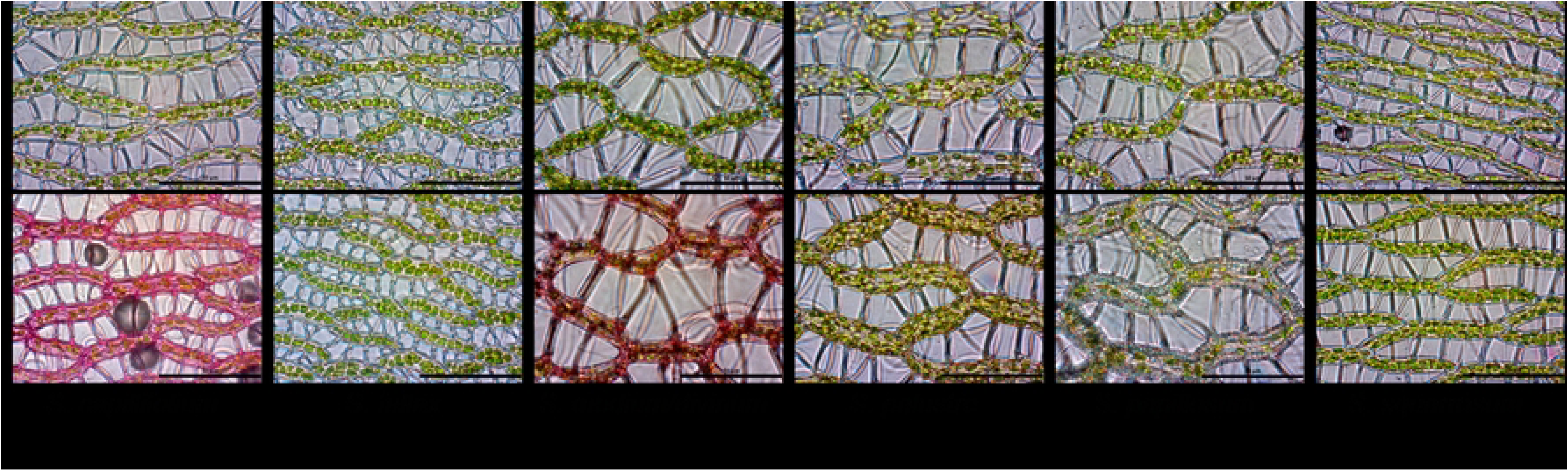
Examples of *Sphagnum* samples at 1000x magnification. Micropropagated (top of pair) and wild-sourced *Sphagnum* samples. Scale bar = 50 µm.

**Table 4.** Comparison of microscopic features between micropropagated and wild-sourced.

| *Sphagnum* species | cell width ($\mu\text{m}$) | | chloroplast no. | |
| --- | --- | --- | --- | --- |
|  | micropropagated | wild-sourced | micropropagated | wild-sourced |
| *S. capillifolium* | $7.14 \pm 0.91$ | $6.65 \pm 0.65$ | $16.6 \pm 2.7^*$ | $10.6 \pm 1.1$ |
| *S. fallax* | $7.14 \pm 0.83$ | $6.96 \pm 0.39$ | $11.1 \pm 2.7$ | $7.1 \pm 1.7$ |
| *S. medium/divinum* | $9.86 \pm 0.72$ | $10.10 \pm 1.36$ | $16.7 \pm 2.6$ | $13.1 \pm 2.9$ |
| *S. palustre* | $12.37 \pm 0.93$ | $11.05 \pm 1.24$ | $24.5 \pm 3.7^*$ | $19.8 \pm 4.3$ |
| *S. papillosum* | $9.88 \pm 0.59$ | $9.84 \pm 0.31$ | $25.6 \pm 3.9^{**}$ | $12.2 \pm 2.2$ |
| *S. squarrosum* | $5.65 \pm 0.59^*$ | $7.30 \pm 1.36$ | $11.9 \pm 2.4$ | $15.1 \pm 3.1$ |

Mean chlorocyst (cell) width ( $\mu\text{m}$ ) and mean number of chloroplasts per chlorocyst; mean values ( $n = 9$ ) ± SD; significant differences (ANOVA Tukey post-hoc HSD) between each pair are indicated: $*p < 0.05$ ; $**p < 0.001$ .

## Discussion

The routine micropropagation of *Sphagnum* in controlled environments has enabled a comparison of species physiology when grown under the same conditions without the confounding influences of habitat conditions and plastic responses. The maximum photosynthesis (P_max_) capacity and respiration rate measured on a range of *Sphagnum* species were broadly in line with the published data for these species (S1 Table), although rates varied more across wild-sourced than micropropagated species, and all were typically much lower than vascular plants [8,11,30,38,39,40].

Several authors [8,9,40] suggest a rank in *Sphagnum* species photosynthesis rates, corresponding generally to the accepted phylogenetic order of growth rates and a gradient in production depending on light, moisture and nutrient levels [39]. In general, highly productive *Sphagna* tend to be competitive or ruderal [41] species of hollows (such as *S. squarrosum*, *S. fallax* and *S. fimbriatum*), thriving in shaded, high moisture and nutrient environments (i.e., low-stress), and adopting an open, loose growth habit [9]. This open, loose growth habit also allows more of the plant access to any available light for photosynthesis and faster growth rates [42]. These species also tend to be green, likely due to a high chlorophyll content [9] with strong shoot growth and large capitula [29]. *Sphagna* with lower productivity tend to be stress-adapted species (such as *S. capillifolium*, *S. medium/divinum* and *S. papillosum*), which grow in open (unshaded), ombrotrophic bogs, subject to high light intensity, occasional drought, and low nutrient levels (i.e., high-stress) [10,11]. In these conditions, short, dense growth forms for greater acquisition and retention of moisture [9,23] and perhaps tolerance of photoinhibition become more important to survival than the capacity for rapid growth [8,23]. Other more generalist species (such as *S. palustre*, and some other loose hummock-forming species) thrive at points along this stress gradient. There are few *Sphagna* in unshaded, dry sites. Thus, capacity for photosynthesis is driven by moisture, light and nutrient levels, and different phylogenetic traits, as described above, give *Sphagnum* species competitive advantages to thrive in particular habitats [25]. This is reflected in our study, and to answer research question 1), whether grown in the same conditions (micropropagated) or sourced from the wild the species with the highest capacity for photosynthesis and respiration (on a dry matter basis) was *S. squarrosum* and the lowest was *S. medium/divinum*, suggesting that the accepted rank, as described above, does have a phylogenetic basis.

The wide range of P_max_ rates across samples in this study was likely due to the diversity of *Sphagnum* species and sources: micropropagated species grown in a commercial greenhouse environment (although original source material was from different sites); wild-sourced species developed each with a range of nutrient and shade regimes in their habitat. It is notable that *S. medium/divinum* P_max_ rates were higher in shaded than open sites in a study by Bengtsson *et al*. [40] (S1 Table). This not only demonstrates the plasticity of this species in its adaptation to a range of environmental conditions but suggests that a shaded environment is likely to promote moisture retention in *Sphagnum* due to reduced evaporative effects of wind and heat, which subsequently supports photosynthesis [9]. Regimes for storage, hydration, acclimatisation and analysis of samples were standardised in our study. Hájek [25] suggested the water content of ‘well-hydrated’ *Sphagnum* is 1500 to 3000% and both micropropagated and wild-sourced samples in this study were within this range. However, wild-sourced samples were at the lower end of the range and had 66% of the water content of micropropagated samples, which may have had some influence on comparative photosynthesis rates.

Micropropagated *Sphagnum* species had a markedly greater response to increasing light levels and higher P_max_ and respiration rates than wild-sourced, on a dry weight basis, which supports research question 2) that there is a difference in photosynthetic capacity and respiration between micropropagated and wild-sourced *Sphagnum* species. Across all species samples, the wild-sourced *Sphagnum* P_max_ was 34.8% (by dry weight) that of the micropropagated *Sphagnum*. This is perhaps because the micropropagated samples in a shaded commercial greenhouse had not yet developed photo-inhibitive adaptations to light stress. However, in both micropropagated and wild-sourced samples, the competitive, shade species, *S. squarrosum* had the highest rates of P_max_, and respiration and the lowest bulk density (mass/volume), and the stress-adapted species, *S. medium/divinum* had the lowest rates of P_max_ and respiration, and the highest bulk density, demonstrating some parity in species behaviour, despite the tissue-culture process.

P_max_ and respiration rates were positively related throughout as typically found across ecosystems [43], most strongly in wild-sourced species suggesting this more mature, natural material may have reached an equilibrium within its habitat, compared with the more even photosynthetic response of micropropagated species to light. There was a higher ratio of P_max_ to respiration in micropropagated than in wild-sourced species, despite generally higher micropropagated respiration rates, suggesting potential carbon balance benefits on application to the field, at least in the early stages of establishment.

Differences in P_max_ between micropropagated and wild-sourced samples were particularly apparent in species from Section *Sphagnum*: *S. medium/divinum, S, palustre* and *S. papillosum*, where photosynthetic activity was noticeably lower in wild-sourced than micropropagated samples. The difference between P_max_ rates of micropropagated and wild-sourced samples of *S. capillifolium* was not so marked as other species. This species is adapted to a range of peatland habitats and ecological niches, and has tolerance to shade [10]. Wild-sourced, shade-tolerant species, *S. squarrosum, S. fallax* and *S. palustre*, had the highest P_max_ levels at the lowest light levels, on a dry weight basis, which concurs with findings by Rice *et al.* [8] and Hájek *et al*. [9].

The DW bulk density of wild-sourced *Sphagnum* for each species studied was significantly greater than that of micropropagated species, which were established under favourable light and moisture regimes, and were also in the early stages of rapid, linear growth [11]. There was also a relationship between increasing bulk density and declining P_max_ in wild-sourced *Sphagnum* samples but not in micropropagated samples. In wild-sourced samples, there were two distinct groups by dry weight bulk density*: S. fallax, S. palustre*, *S. squarrosum* (low bulk density), and *S. capillifolium, S. papillosum*, *S. medium/divinum* (high bulk density), showing obvious differences between shade-adapted, moisture-dependent species, and light-adapted, moisture-retaining species, which are only fully expressed in the natural environment.

Micropropagated *Sphagnum* species were not visually different to those wild-sourced, although some characteristics, such as *S. papillosum* cell papillae, were less developed in micropropagated samples (Fig 8), and there was less colour expression in micropropagated *Sphagnum* (Fig 1) due to lack of exposure to stress conditions. Chlorocyst size was not significantly different despite the relative immaturity of micropropagated samples, apart from *S. squarrosum* species where those of wild-sourced samples were significantly larger than micropropagated samples and the number of chloroplasts per cell was also greater. *S. squarrosum* samples were sourced from a nutrient-rich, shaded environment, beneficial to continued upward growth; greater cell size and number of chloroplasts were likely based on maturity and optimum growing conditions. In all other species, there was a significantly greater number of chloroplasts in micropropagated than in wild-sourced samples, apart from *S. palustre*, where the difference was not statistically significant. This is consistent with micropropagated plants being in the early stages of rapid growth [11] but also not exposed to conditions of high light intensity and low moisture, and so they were perhaps acting more like shade plants [44]. Therefore, the aspect of research question 3) examining whether any differences in chlorocyst size and chloroplast number between micropropagated and wild-sourced *Sphagnum* species may affect their photosynthesis and respiration did not lead to a clear conclusion, as any differences were not directly linked to different photosynthetic capacity. However, the high rates of photosynthesis in micropropagated *Sphagnum*, the greater number of capitula per sample, as well as similar features to wild-sourced, suggest that successful establishment in optimum field conditions is likely.

Nutrient analysis of samples allowed further examination of causes of difference in photosynthesis rates. micropropagated samples contained significantly higher levels of macronutrients, essential for plant photosynthesis and growth, than wild-sourced samples, and as CO_2_ uptake and emission rates were also higher in micropropagated *Sphagnum* it appears that the aspect of research question 3) which examined whether any differences in nutrient content between micropropagated and wild-sourced *Sphagnum* species may affect their photosynthesis and respiration, was supported.

Higher levels of macronutrients in micropropagated *Sphagnum* may be explained through standard horticultural processes of nutrient application associated with micropropagated production. Conversely, wild-sourced *Sphagnum* had higher levels than micropropagated of micronutrients, which are a smaller component of plant nutritional requirements but also support plant health. Concentrations of micronutrients may have become diluted in micropropagated plants due to their rapid growth rate. Additionally, higher levels of elements considered detrimental to plant health (such as Al, Fe and Pb) in some wild-sourced species may reflect the capability of bryophytes to absorb certain levels of pollution and act as bioindicators [4,45].

The three main macronutrients associated with plant growth and photosynthesis, N, P and K, were strongly positively associated with P_max_ overall in this study and reflected species’ differences in macronutrient uptake and P_max_ rates. For example, wild-sourced species with the highest content of NPK were *S. squarrosum* and *S. palustre,* and those with the lowest content were *S. papillosum* and *S. medium/divinum*, which corresponded with P_max_ rates for those species.

In vascular plants, there is a strong positive relationship between both net photosynthesis and dark respiration and leaf N content in a wide range of global biomes [38]. Typically, mosses translocate nutrients into their tissues gradually over the growing season [46] and the N:P and N:K ratios remain balanced [47]. Granath *et al*. [48] and Mazziotta *et al*. [27] found a positive relationship between N concentration and photosynthesis rates in *Sphagnum* (as in this study). However, *Sphagnum* growth reportedly improves with increasing levels of N until a ‘critical concentration’ is reached, at which there is no further promotion of plant growth despite increasing levels of N in tissues [49]. In the context of aerial nitrogen pollution, Press *et al*. [50] found that increasing inorganic nitrogen supply slowed *Sphagnum* growth. Saturated N levels limit further P- and K-accumulation, and so growth becomes P- and K-limited [4,51,52]. Excess N is also leached into the environment, promoting the growth of vascular plants which may outcompete *Sphagnum* in the field [26,52,53].

The balance between N, P and K levels in *Sphagnum* tissues appears to be similar throughout natural bog settings, but differences may occur due to growth habit and conditions. Wang and Moore [54] reported averages of N = 9, P = 0.55, K = 7.5 mg g^-1^ DW content for ‘hummock’ *Sphagna* (*S. capillifolium* and *S. medium/divinum*) at the Mer Bleue ombrotrophic bog in Canada. Bragazza *et al*. [4] had similar values of N = 8.2 and 9.2, P = 0.41 and 0.44, K = 4 and 4.44 mg g^-1^ DW for hummock and lawn species respectively in European mires, which are reported to have higher N deposition than Canadian mires. NPK levels of wild-sourced species in this study had higher N values but were otherwise similar to European values: N = 12.0 ± 2.13, P = 0.44 ± 0.24, K = 4.05 ± 1.87 mg g^-1^ DW. Wild-sourced *S. squarrosum*, however, had a higher NPK content (N = 20.9 ± 2.05, P = 1.83 ± 0.16, K = 13.6 ± 0.75 mg g^-1^ DW) and, as the sample was sourced from wet woodland, suggests that nutrients may continue to accumulate in fast-growing *Sphagnum* in conditions of shade [55] and optimum moisture [33].

All micropropagated species had a high NPK content (N = 24.2 ± 4.41, P = 2.55 ± 0.49, K = 13.2 ± 2.79 mg g^-1^ DW) due to greenhouse nutrient additions. *Sphagnum* N concentration thresholds of 11 to 12 mg g^-1^ [4,52], 15 mg g^-1^ (for *S. recurvum* - Section *Cuspidata*) [56] and 20 mg g^-1^ (for a range of hummock and hollow species) [53] have been reported. Nitrogen content in wild-sourced *S. squarrosum* in this study was at the top of the reported threshold range, with no apparent limitation of P or K, although this may be due to the fast-growing nature of this species, and nutrients are readily replenished by the woodland habitat from which it was sourced. Nitrogen content in micropropagated samples was well above the highest reported threshold – as high as 30 mg g^-1^ in some samples, with no evidence of toxicity or limitation of P or K.

Growth reduction is reported when *Sphagnum* experiences a considerable increase in N availability relative to conditions at the site from which it was obtained [57], implying that *Sphagnum* tolerance of N is site-adapted. Bragazza *et al*. [4] found that, at elevated N levels the N:P ratio increased to 33.8 and 33.6 and N:K ratio increased to 3.7 and 4.0 for hummock and lawn species respectively. In this study, there were very high N:P and marginally high N:K ratios in *S. medium/divinum* (64.6 and 5.24) and *S. papillosum* (42.6 and 3.86 respectively) from open, ombrotrophic bogs compared to N:P and N:K ratios in *S. squarrosum* (11.4 and 1.54) and *S. palustre* (17.5 and 2.05 respectively) (from higher-nutrient, shaded environments) which are more aligned to ratios found in low nutrient sites by Wang and Moore [54] and Bragazza *et al*. [4]. In wild-sourced *Sphagnum* in this study, the lowest P_max_ was found in *S. papillosum* and *S. medium/divinum*, and the highest in *S. squarrosum* and *S. palustre*.

Although the plants with a high N:P ratio (*S. papillosum* and *S. medium/divinum*) appeared healthy and are likely adapted to these particular levels of nutrients, limitation of P by higher levels of N (as described above) may have reduced their photosynthetic capacity [58], although Granath *et al*. [48] found that the negative effect of limited P was on production rather than photosynthetic rate.

This difference in stoichiometry between micropropagated and wild-sourced plants may be due both to issues of maturity [47], as levels of N may be naturally higher in young than in mature plants [49], and site adaptation in wild-sourced samples [57] i.e., plant adaptations to environmental stressors inhibited photosynthetic potential in wild-sourced plants. Although micropropagated plants were ‘mature’ in terms of development, they could be classed as ‘young’, being grown directly from tissue culture to the point at which they are available as BeadaHumok™ ‘plugs’ after only several months of growth. Micropropagated plants do not appear to be P- or K-limited (N:P = 9.61 ± 1.24 and N:K = 1.85 ± 0.09) suggesting that the right balance of nutrients is being applied in the horticultural process for these new plants.

The most important aspect of *Sphagnum* reintroduction to peatland restoration sites is rapid establishment and growth, particularly lateral growth, to make an intact carpet as quickly as possible [59]. This will progress restoration by keeping a cool, moist layer at the peat surface [60], reducing evaporation and encouraging development of an acrotelm to promote peat accumulation [18,61,62,63]. The capacity of micropropagated *Sphagnum* to absorb Nitrogen above normally limited thresholds may also be useful on application to ex-agricultural sites as a paludiculture crop or for restoration purposes, at least in the critical establishment phase. This study on differences between micropropagated and wild-sourced *Sphagnum* in terms of net photosynthesis, phylogenetic and adaptive traits, and nutrient assimilation highlights potential reasons for the successful establishment and growth of micropropagated products reported in large-scale restoration projects [19,20,21] and their potential contribution to CO_2_ uptake. Future work could usefully determine whether the advantages conveyed in these early growth stages of micropropagated *Sphagnum* moss persist over longer time periods in the wild.

## Acknowledgements

This paper was derived from a doctoral thesis (PhD) part-funded by Micropropagation Services (EM) Ltd, match-funded by Manchester Metropolitan University (MMU). The authors wish to thank David McKendry and technical officers at MMU for their support, and are grateful to the examiners of the PhD thesis who provided valuable comments and corrections.

## Supporting information

**S1 Table. Comparison of light-saturated photosynthesis and respiration rates between samples in this study and values in the literature.** Light-saturated photosynthesis (P_max_) and respiration (Resp) rates are expressed by dry weight (nmol CO_2_ g^-1^ s^-1^); leaf gas exchange sign convention used (i.e. CO_2_ uptake positive and CO_2_ emission negative); Bengtsson *et al*., 2016 values estimated from graphs; C = competitive; R = ruderal; S = stress-tolerant (from Kangas *et al*., 2014 [sensu 41]); *S. magellanicum* reported assumed to be *S. medium/divinum*. **\***Vascular leaves range from slow growing evergreens to fast growing herbs.

## References

1. Verhoeven JT, Liefveld WM. The ecological significance of organochemical compounds in Sphagnum. Acta Botanica Neerlandica. 1997 Jan 1;46(2):117–30.

2. Rydin H, Jeglum JK, Bennett KD. The biology of peatlands, 2e. OUP Oxford; 2013 Jul 18.

3. Lindsay R, Birnie R, Clough J. Peat bog ecosystems: key definitions. International Union for the Conservation of Nature; 2014 Nov 5. [Cited 2024 Apr 24]. Available from: https://repository.uel.ac.uk/item/85870

4. Bragazza L, Tahvanainen T, Kutnar L, Rydin H, Limpens J, Hájek M, Grosvernier P, Hájek T, Hajkova P, Hansen I, Iacumin P. Nutritional constraints in ombrotrophic Sphagnum plants under increasing atmospheric nitrogen deposition in Europe. New Phytologist. 2004 Sep 1:609–16.

5. Fritz C, Lamers LP, Riaz M, van den Berg LJ, Elzenga TJ. Sphagnum mosses-masters of efficient N-uptake while avoiding intoxication. PloS one. 2014 Jan 9;9(1):e79991.

6. van Breemen N. How Sphagnum bogs down other plants. Trends in ecology & evolution. 1995 Jul 1;10(7):270–5.

7. Malmer N, Albinsson C, Svensson BM, Wallén B. Interferences between Sphagnum and vascular plants: effects on plant community structure and peat formation. Oikos. 2003 Mar;100(3):469–82.

8. Rice SK, Aclander L, Hanson DT. Do bryophyte shoot systems function like vascular plant leaves or canopies? Functional trait relationships in Sphagnum mosses (Sphagnaceae). American Journal of Botany. 2008 Nov;95(11):1366–74.

9. Hájek T, Tuittila ES, Ilomets M, Laiho R. Light responses of mire mosses–a key to survival after water-level drawdown? Oikos. 2009 Feb;118(2):240–50.

10. Bonnett SA, Ostle N, Freeman C. Short-term effect of deep shade and enhanced nitrogen supply on *Sphagnum capillifolium* morphophysiology. Plant Ecology. 2010 Apr;207:347–58.

11. Laine AM, Juurola E, Hájek T, Tuittila ES. Sphagnum growth and ecophysiology during mire succession. Oecologia. 2011 Dec;167:1115–25.

12. Joosten H, Sirin A, Couwenberg J, Laine J, Smith P. The role of peatlands in climate regulation. In: Peatland Restoration and Ecosystem Services: Science, Policy and Practice 2016 Jun 23 (pp. 63-76). Cambridge, UK: Cambridge University Press.

13. Gregg R, Elias J, Alonso I, Crosher I, Muto P, Morecroft M. Carbon sequestration by habitat: a review of the evidence 2021. Natural England Research Report NERR094. Natural England, York.

14. Evans C, Artz R, Moxley J, Smyth MA, Taylor E, Archer E, Burden A, Williamson J, Donnelly D, Thomson A, Buys G. Implementation of an emissions inventory for UK peatlands. Centre for Ecology and Hydrology; 2017 Dec 20.

15. BEIS (2019) 2017 UK Greenhouse Gas emissions, final figures. Statistical Release: National Statistics. Department for Business, Energy and Industrial Strategy. 2019 Feb 5 [Cited 2024 April 24]. Available from: https://assets.publishing.service.gov.uk/media/5c584a12e5274a317cc0eaf7/2017_Final_emissions_statistics_-_report.pdf

16. Wilson D, Farrell C, Mueller C, Hepp S, Renou-Wilson F. Rewetted industrial cutaway peatlands in western Ireland: a prime location for climate change mitigation? Mires & Peat. 2013 Jan 1;11.

17. Renou-Wilson F, Moser G, Fallon D, Farrell CA, Müller C, Wilson D. Rewetting degraded peatlands for climate and biodiversity benefits: Results from two raised bogs. Ecological Engineering. 2019 Feb 1;127:547–60.

18. Quinty F, Rochefort L. Peatland restoration guide 2003. [Cited 2024 May 27]. Available from: https://www1.up.poznan.pl/glinbar/wp-content/uploads/2015/03/Restoration-Guide_2nd_2003.pdf

19. Caporn SJ, Rosenburgh AE, Keightley AT, Hinde SL, Riggs JL, Buckler M, Wright NA. Sphagnum restoration on degraded blanket and raised bogs in the UK using micropropagated source material: a review of progress Mires & Peat. 2018 May 11;20.

20. Crouch T. Kinder Scout Sphagnum Trials: 2018 Update Report, Moors for the Future Report, Edale. [Cited 2024 May 27]. Available from: https://www.moorsforthefuture.org.uk/data/assets/pdf_file/0027/93933/MFFP-Kinder-Scout-Sphagnum-Trials-Update-Report-2018.pdf

21. Pilkington M, Walker J, Fry C, Eades P, Meade R, Pollett N, Rogers T, Helliwell T, Chandler D, Fawcett E, Keatley T. Diversification of Molinia-dominated blanket bogs using Sphagnum propagules. Ecological Solutions and Evidence. 2021 Oct;2(4):e12113.

22. Bengtsson F, Rydin H, Baltzer JL, Bragazza L, Bu ZJ, Caporn SJ, Dorrepaal E, Flatberg KI, Galanina O, Gałka M, Ganeva A. Environmental drivers of Sphagnum growth in peatlands across the Holarctic region. Journal of Ecology. 2021 Jan;109(1):417–31.

23. Loisel J, Gallego-Sala AV, Yu Z. Global-scale pattern of peatland Sphagnum growth driven by photosynthetically active radiation and growing season length. Biogeosciences. 2012 Jul 30;9(7):2737–46.

24. Hassel K, Kyrkjeeide MO, Yousefi N, Prestø T, Stenøien HK, Shaw JA, Flatberg KI. Sphagnum divinum (sp. nov.) and S. medium Limpr. and their relationship to S. magellanicum Brid. Journal of Bryology. 2018 Jul 3;40(3):197–222.

25. Hájek T. Physiological ecology of peatland bryophytes. In Photosynthesis in bryophytes and early land plants 2013 Aug 7 (pp. 233–252). Dordrecht: Springer Netherlands.

26. Bubier JL, Moore TR, Bledzki LA. Effects of nutrient addition on vegetation and carbon cycling in an ombrotrophic bog. Global Change Biology. 2007 Jun;13(6):1168–86.

27. Mazziotta A, Granath G, Rydin H, Bengtsson F, Norberg J. Scaling functional traits to ecosystem processes: Towards a mechanistic understanding in peat mosses. Journal of Ecology. 2019 Mar;107(2):843–59.

28. Aldous AR. Nitrogen translocation in Sphagnum mosses: effects of atmospheric nitrogen deposition. New Phytologist. 2002 Nov;156(2):241–53.

29. Laing C, Granath G, Belyea LR, Allton KE, Rydin H. Tradeoffs and scaling of functional traits in Sphagnum as drivers of carbon cycling in peatlands. Oikos. 2014 Jul;123(7):817–28.

30. Haraguchi A, Yamada N. Temperature dependency of photosynthesis of Sphagnum spp. distributed in the warm-temperate and the cool-temperate mires of Japan. American Journal of Plant Sciences. 2011 Nov 25;2(05):716.

31. Dossa GG, Paudel E, Wang H, Cao K, Schaefer D, Harrison RD. Correct calculation of CO_2_ efflux using a closed-chamber linked to a non-dispersive infrared gas analyzer. Methods in Ecology and Evolution. 2015 Dec;6(12):1435–42.

32. Limpens J, Berendse F. How litter quality affects mass loss and N loss from decomposing Sphagnum. Oikos. 2003 Dec;103(3):537–47.

33. McNeil P, Waddington JM. Moisture controls on Sphagnum growth and CO_2_ exchange on a cutover bog. Journal of applied ecology. 2003 Apr;40(2):354–67.

34. Smith AJ. The moss flora of Britain and Ireland, 2e. Cambridge university press; 2004 Sep 23.

35. Laine J, Flatberg KI, Harju P, Timonen T, Minkkinen KJ, Laine A, Tuittila ES, Vasander HT. Sphagnum mosses: the stars of European mires. Sphagna Ky.; 2018 Aug 31.

36. Atherton I, Bosanquet S, Lawley M, editors. Mosses and liverworts of Britain and Ireland: a field guide. Plymouth: British Bryological Society; 2010.

37. Hammer Ø, Harper DA. Past: paleontological statistics software package for education and data analysis. Palaeontologia electronica. 2001;4(1):1.

38. Reich PB, Walters MB, Ellsworth DS. From tropics to tundra: global convergence in plant functioning. Proceedings of the National Academy of Sciences. 1997 Dec 9;94(25):13730–4.

39. Kangas L, Maanavilja L, Hájek T, Juurola E, Chimner RA, Mehtätalo L, Tuittila ES. Photosynthetic traits of Sphagnum and feather moss species in undrained, drained and rewetted boreal spruce swamp forests. Ecology and Evolution. 2014 Feb;4(4):381–96.

40. Bengtsson F, Granath G, Rydin H. Photosynthesis, growth, and decay traits in Sphagnum–a multispecies comparison. Ecology and Evolution. 2016 May;6(10):3325–41.

41. Grime JP. Evidence for the existence of three primary strategies in plants and its relevance to ecological and evolutionary theory. The american naturalist. 1977 Nov 1;111(982):1169–94.

42. Krebs M, Gaudig G, Joosten H. Record growth of Sphagnum papillosum in Georgia (Transcaucasus): rain frequency, temperature and microhabitat as key drivers in natural bogs. Mires & Peat. 2016 Jul 1;18.

43. Chapin FS, Matson PA, Vitousek PM, Chapin FS, Matson PA, Vitousek PM. Principles of terrestrial ecosystem ecology. 2011.

44. Marschall M, Proctor MC. Are bryophytes shade plants? Photosynthetic light responses and proportions of chlorophyll a, chlorophyll b and total carotenoids. Annals of botany. 2004 Oct 1;94(4):593–603.

45. Blagnytė R, Paliulis D. Research into heavy metals pollution of atmosphere applying moss as bioindicator: a literature review. Environmental Research, Engineering and Management. 2010 Dec 21;54(4):26–33.

46. Chapin III FS, Johnson DA, McKendrick JD. Seasonal movement of nutrients in plants of differing growth form in an Alaskan tundra ecosystem: implications for herbivory. The Journal of Ecology. 1980 Mar 1:189–209.

47. Sterner RW, Elser JJ. Ecological stoichiometry: the biology of elements from molecules to the biosphere. In Ecological stoichiometry 2017 Feb 15. Princeton university press.

48. Granath G, Strengbom J, Rydin H. Direct physiological effects of nitrogen on Sphagnum: a greenhouse experiment. Functional Ecology. 2012 Apr;26(2):353–64.

49. Barker AV, Pilbeam DJ, editors. Handbook of plant nutrition. CRC press; 2015.

50. Press MC, Woodin SJ, Lee JA. The potential importance of an increased atmospheric nitrogen supply to the growth of ombrotrophic Sphagnum species. New Phytologist. 1986 May;103(1):45–55.

51. Aerts R, Wallen BO, Malmer N. Growth-limiting nutrients in Sphagnum-dominated bogs subject to low and high atmospheric nitrogen supply. Journal of ecology. 1992 Mar 1:131–40.

52. Lamers LP, Bobbink R, Roelofs JG. Natural nitrogen filter fails in polluted raised bogs. Global Change Biology. 2000 Jun;6(5):583–6.

53. Berendse F, Van Breemen N, Rydin H, Buttler A, Heijmans M, Hoosbeek MR, Lee JA, Mitchell E, Saarinen T, Vasander H, Wallén B. Raised atmospheric CO_2_ levels and increased N deposition cause shifts in plant species composition and production in *Sphagnum* bogs. Global change biology. 2001 May;7(5):591–8.

54. Wang M, Moore TR. Carbon, nitrogen, phosphorus, and potassium stoichiometry in an ombrotrophic peatland reflects plant functional type. Ecosystems. 2014 Jun;17:673–84.

55. Ma JZ, Bu ZJ, Zheng XX, Ge JL, Wang SZ. Effects of shading on relative competitive advantage of three species of Sphagnum. Mires & Peat. 2015 Jun 1;16.

56. van Der Heijden E, Verbeek SK, Kuiper PJ. Elevated atmospheric CO_2_ and increased nitrogen deposition: effects on C and N metabolism and growth of the peat moss Sphagnum recurvum P. Beauv. var. mucronatum (Russ.) Warnst. Global Change Biology. 2000 Feb;6(2):201–12.

57. Limpens J, Berendse F. Growth reduction of Sphagnum magellanicum subjected to high nitrogen deposition: the role of amino acid nitrogen concentration. Oecologia. 2003 May;135:339–45.

58. Reich PB, Oleksyn J, Wright IJ. Leaf phosphorus influences the photosynthesis–nitrogen relation: a cross-biome analysis of 314 species. Oecologia. 2009 May;160:207–12.

59. Rochefort L. Sphagnum—a keystone genus in habitat restoration. The Bryologist. 2000 Sep;103(3):503–8.

60. Waddington JM, Warner K. Atmospheric CO_2_ sequestration in restored mined peatlands. Ecoscience. 2001 Jan 1;8(3):359–68.

61. Lucchese M, Waddington JM, Poulin M, Pouliot R, Rochefort L, Strack M. Organic matter accumulation in a restored peatland: Evaluating restoration success. Ecological Engineering. 2010 Apr 1;36(4):482–8.

62. Waddington JM, Lucchese MC, Duval TP. Sphagnum moss moisture retention following the re-vegetation of degraded peatlands. Ecohydrology. 2011 May;4(3):359–66.

63. Worrall F, Chapman P, Holden J, Evans C, Artz R, Smith P, Grayson R. A review of current evidence on carbon fluxes and greenhouse gas emissions from UK peatlands.

